# Contribution of bacterial and host factors to pathogen “blooming” in a gnotobiotic mouse model for *Salmonella enterica* serovar Typhimurium-induced enterocolitis

**DOI:** 10.1101/2023.08.21.554185

**Authors:** Markus Beutler, Claudia Eberl, Debora Garzetti, Simone Herp, Philipp Münch, Diana Ring, Tamas Dolowschiak, Sandrine Brugiroux, Patrick Schiller, Saib Hussain, Marijana Basic, Andre Bleich, Barbel Stecher

## Abstract

Inflammation has a pronounced impact on the intestinal ecosystem by driving an expansion of facultative anaerobic bacteria at the cost of obligate anaerobic microbiota. This pathogen “blooming” is also a hallmark of enteric *Salmonella* enterica serovar Typhimurium (*S*. Tm) infection. Here, we analyzed the contribution of bacterial and host factors to *S*. Tm “blooming” in a gnotobiotic mouse model for *S.* Tm-induced enterocolitis. Mice colonized with the Oligo-Mouse-Microbiota (OMM^12^), a minimal bacterial community, develop fulminant colitis by day 4 after oral infection with wild type *S*. Tm but not with an avirulent mutant. Inflammation leads to pronounced reduction in overall intestinal bacterial loads, distinct microbial community shifts and pathogen blooming (relative abundance >50%). *S.* Tm mutants attenuated in inducing gut inflammation generally elicit less pronounced microbiota shifts and reduction in total bacterial loads. In contrast, *S.* Tm mutants in nitrate respiration, salmochelin production and ethanolamine utilization induced strong inflammation and *S*. Tm “blooming”. Therefore, individual *Salmonella*-specific inflammation-fitness factors seem to be of minor importance for competition against this minimal microbiota in the inflamed gut. Finally, we show that antibody-mediated neutrophil depletion normalized gut microbiota loads but not intestinal inflammation or microbiota shifts. This suggests that neutrophils equally reduce pathogen and commensal bacterial loads in the inflamed gut.

## Introduction

*Salmonella enterica* serovar Typhimurium (*S*. Tm) is a frequent cause of Salmonellosis in humans with millions of infections worldwide (1). The infection is usually self-limiting however, in very young, old or immunocompromised patients *S.* Tm can induce life threatening systemic disease with several thousands of deaths per year (2). The intestinal microbiota efficiently lowers the risk of infection by mediating colonization resistance (CR). When CR is reduced such as upon antibiotic therapy (3), a high-fat meal (4) or an infant or low-complex microbiota (5), *S.* Tm can invade the gut ecosystem and cause disease. In mice, the infection proceeds in several phases (6): First, *S.* Tm expands to high numbers in the gut by consuming hydrogen (7), diet- and microbiota-liberated mucosal carbohydrates (8, 9), oxygen (10), anaerobic electron acceptors and fumarate (11, 12). Only after reaching a certain density (>10^5^ cfu/g), *S.* Tm invades the gut mucosa at sufficient numbers and triggers intestinal inflammation (13, 14).

Gut inflammation dramatically alters the intestinal environment, leading to drastic changes in microbial community composition and favoring the *S.* Tm overgrowth (*Salmonella* “blooming”) over obligate anaerobic commensal bacteria (15, 16). Evidently, *Salmonella* “blooming” is caused by two contrasting effects of inflammation (17, 18): (i) selective fostering of pathogen growth by nutritional changes and (ii) differential killing of the mostly anaerobic commensal microbiota by innate immune defense mechanisms against which *Salmonella* is more resistant. In this study, we analyzed the contribution of both, bacterial and host factors to *Salmonella* “blooming” in a gnotobiotic mouse model for *S.* Tm-induced enterocolitis.

Increase in the concentration of luminal oxygen levels well as anaerobic electron acceptors generated by the inflamed mucosa (10, 19) favor *S.* Tm growth over exclusively fermenting anaerobes (11, 20). This enables the pathogen to consume host-, diet and microbiota-derived metabolites including mucin-derived sugars, ethanolamine, 1,2-propanediol, fructose-asparagine, succinate and lactate (21–26) and gain a competitive advantage over competing commensals. Production of the siderophore salmochelin, a glycosylated variant of enterochelin which is not bound by the antimicrobial protein lipocalin 2(LCN-2), also boosts *S.* Tm in the inflamed gut (27). On the other hand, acute *S.* Tm inflammation also exerts significant “collateral damage” to the beneficial microbiota (28). The microbiota is inhibited by antimicrobial molecules (27, 29) and bile salts (30), against which *Salmonella* exhibits a higher degree of resistance.

Moreover, neutrophils which are a hallmark of acute *Salmonella*-induced inflammation (31, 32), infiltrate the mucosa where they effectively reduce pathogen tissue loads by an NADPH-oxidase-mediated defense (13, 33). Transmigrated live neutrophils, which engulf luminal S. Tm, are present in the gut lumen early after infection (34). Neutrophils also form intraluminal structures (“casts” or pseudomembranes), which encapsulate commensals and thus prevent their epithelial contact and translocation (35). In *S.* Tm infection, luminal neutrophils impose a drastic bottleneck upon the pathogen population (36) and are likely to also kill commensals. In particular, the neutrophil effector molecules lipocalin-2, calprotectin and elastase have been associated with microbiota alterations in response to pathogen-induced inflammation in mice (37, 38).

Overall, selective nutritional use and differential killing sustain long-term *S.* Tm “blooming” and the establishment of a “supershedder” state, i.e. individuals that show high pathogen loads in stool (39). However, the relative importance of these contributing effects to *S.* Tm “blooming” remain elusive, as previous studies focused on individual mechanisms and used different experimental protocols and *S.* Tm strains. Moreover, it is challenging to disentangle inflammation- and antibiotic-induced changes in the microbiome in models that rely on antibiotic-pretreatment. To overcome this limitation, we analyzed the course of *S.* Tm-induced inflammation in a gnotobiotic infection model that does not rely on antibiotic-pretreatment. Mice stably colonized with the Oligo-Mouse-Microbiota (OMM^12^; 12 mouse-derived commensals), exhibit partial colonization resistance and develop *S.* Tm gut inflammation after oral infection (40). This model is widely used and steadily gaining ground in research of colonization resistance (4, 9, 41). Here, we provide a detailed characterization of enteric *S.* Tm infection and its effect on the gut microbial community in the OMM^12^ model. We study the course of infection of different *S.* Tm mutants and assess their ability to cause inflammation-induced *S.* Tm “blooming”. Finally, we employ a neutrophil depletion protocol to address the role of neutrophils in *S.* Tm “blooming”, luminal bacterial killing and microbiota alterations.

## Results

### Course of oral *S*. Tm infection in OMM^12^ mice

Mice stably colonized with the OMM^12^ community (OMM^12^ mice) were infected intragastrically (i.g.) with 5 x 10^7^ CFUs of either wild type *S.* Tm (*S*. Tm^WT^) or an avirulent *S.* Tm mutant (*S*. Tm^avir^; Δ*invG*; *sseD*::*aphT*) lacking functional *Salmonella* pathogenicity island (SPI)-1 and SPI-2 encoded type III secretion systems (T3SS). At day 1 to day 4 post infection (p.i.), groups of mice were sacrificed and intestinal and organ samples were taken for analyses (**Fig. 1A**). Cecum loads of both *S.* Tm strains increased steadily from day 1 to day 4 p.i.. No difference was observed between *S*. Tm^avir^ and *S*. Tm^WT^ loads at all tested days (p>0.05, Kruskal-Wallis test with Dunn’s multiple comparison test; **Fig. 1B**). Compared to S. Tm^avir^, *S*. Tm^WT^ loads were higher at day 4 p.i. in the mesenteric lymph nodes (mLN; p<0.01, **Fig. 1C**). No significant difference could be detected in the spleen (p>0.05; **Fig. 1E**). Starting at day 3 p.i., *S*. Tm^WT^ caused strong gut inflammation as determined by vastly increased cecal lipocalin-2 (LCN-2) levels (**Fig. 1F**) and cecal histopathology (**Fig. 1G**). Mice infected with avirulent *S.* Tm did not exhibit overt pathological changes in the cecum or increased LCN-2 (**Fig. 1FG**).

**Fig. 1:**
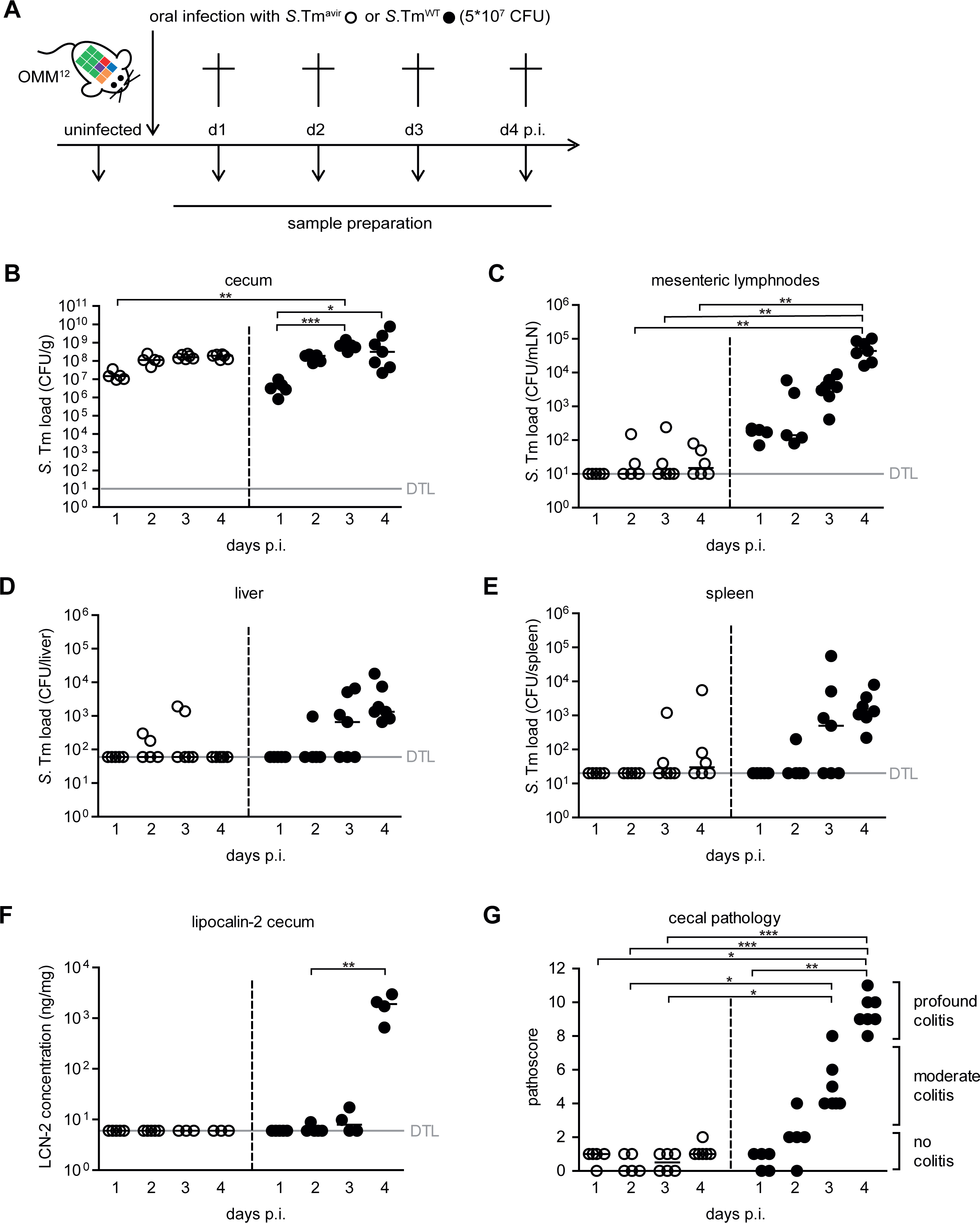
Course of enteric *S*. Tm infection and systemic dissemination. **(A)** Experimental scheme. OMM^12^ mice were orally infected with either *S*. Tm^avir^ or *S*. Tm^WT^. Mice were sacrifices at days 1, 2, 3 and 4 post infection (infection with *S*. Tm^avir^: day 1 p.i. n=5, day 2 p.i. n=5, day 3 p.i. n=6, day 4 p.i. n=6; infection with *S*. Tm^WT^: day 1 p.i. n=5, day 2 p.i. n=5, day 3 p.i. n=7, day 4 p.i. n=7). *S.* Tm loads in **(B)** cecal content, **(C)** mesenteric lymphnodes, **(D)** liver and **(E)** spleen were determined by plating. **(F)** Lipocalin-2 amount in cecal content d4 was determined by ELISA. **(G)** Histopathological analysis of cecal tissue. Cecal tissue sections of the mice were stained with hematoxylin/eosin to determine the degree of submucosal edema, neutrophil infiltration and epithelial damage (1–3: no pathological changes; 4–6: moderate inflammation; above 7: severe inflammation). Statistical analysis was performed using Kruskal-Wallis test with Dunn’s multiple comparison test (* p<0.05, ** p<0.01, *** p<0.001). Each dot represents one mouse, black lines indicate median, grey lines indicate detection limit (DTL).

### *S.* Tm^WT^ infection leads to pathogen blooming, decrease in gut luminal bacterial load and community shifts

Next, we determined OMM^12^ community composition including *S.* Tm loads in cecum content (**Fig. 2, Fig. S1**) and feces (**Fig. S2**) by strain-specific qPCR. Calculations of relative abundance revealed that *S*. Tm^WT^ dominated (rel. abundance >50%) the bacterial community in feces and cecum content by day 4 p.i. (**Fig. 2AC, Fig. S2AC**), coinciding with developing gut inflammation (**Fig. 1FG**). In contrast, no significant increase in relative abundance over time was observed for *S*. Tm^avir^ (**Fig. 2AC, Fig. S2AC**). ADONIS Permanova analysis on the Bray–Curtis distance metric revealed significant differences of microbial community composition by day and between *S*. Tm^WT^ and S. Tm^avir^ infected mice in feces and cecal content (**Fig. 2AB; Fig. S2AB; Table S1; S2**). Strikingly, *S*. Tm^WT^ infection and resulting inflammation led to a drastic decrease in total gut bacterial loads determined by 16SrRNA copies (**Fig. 2D; Fig. S2D**). This was the case for the majority of OMM^12^ strains but *Enterococcus faecalis* KB1 (**Fig. S1**).

**Fig. 2:**
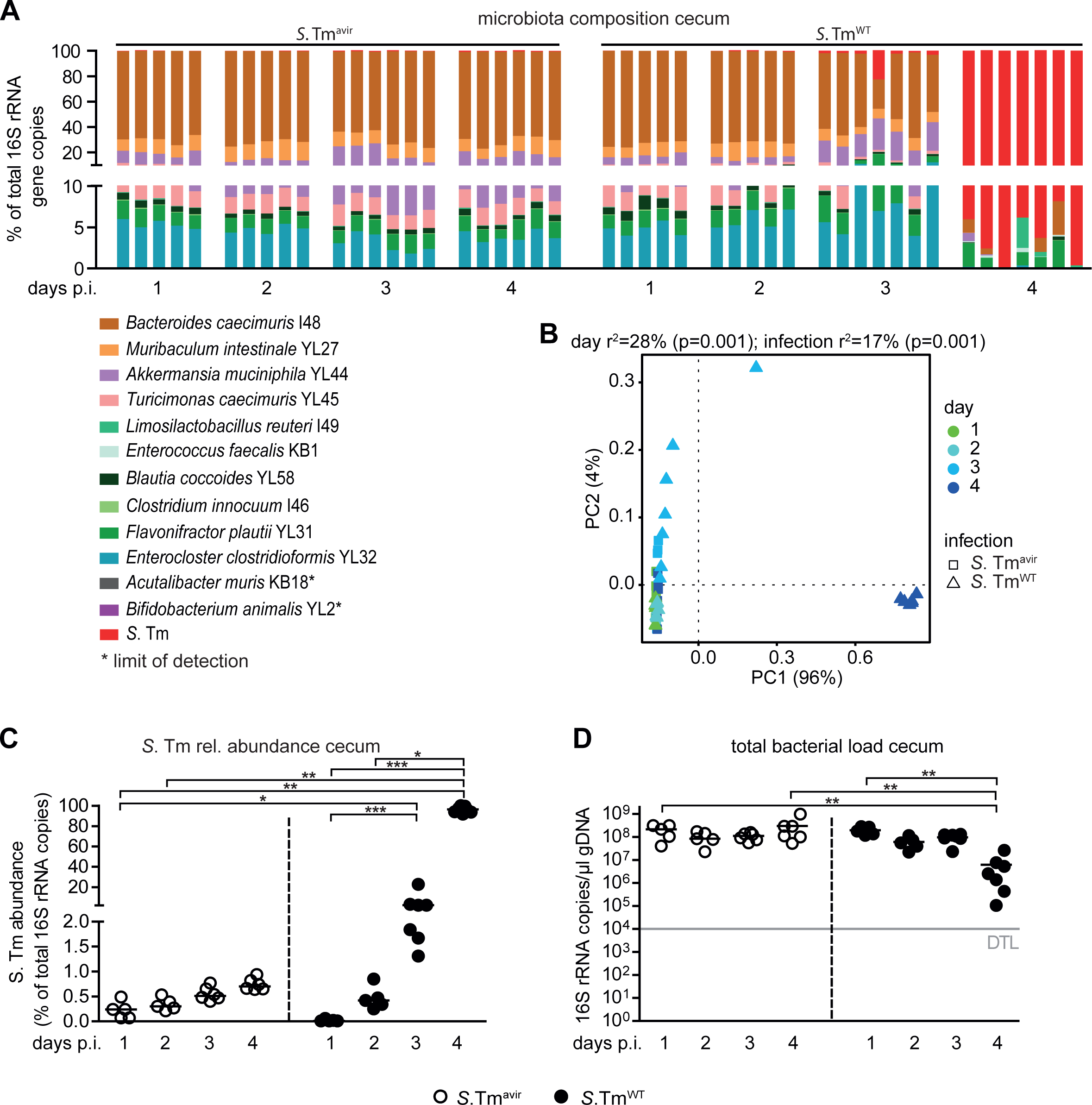
*S*. Tm^WT^ causes pronounced shifts in microbiota composition at day 4 post infection in OMM^12^ mice. **(A)** Analysis of microbiota composition in cecal content. Microbiota composition was determined by strain-specific qPCR assay and is shown as relative abundances of the individual strains (% of cumulated 16S rRNA gene copy numbers). **(B)** PCoA based on the distance matrix of Bray-Curtis dissimilarity of relative OMM^12^ abundance profiles shows the effect of time after infection. Points are colored by time (days) after infection. **(C)** Relative abundance of *S*. Tm in cecal contents at different time points. **(D)** Absolute amount of 16S rRNA gene copies (determined by an universal primer / probe combination). Statistical analysis was performed using Kruskal-Wallis test with Dunn’s multiple comparison test (* p<0.05, ** p<0.01, *** p<0.001). Each dot represents one mouse, black lines indicate median, grey lines indicate detection limit (DTL).

We also infected OMM^12^ mice with *S*. Tm mutants in either the SPI-1 (*S*. Tm^ΔSPI-1^)- or the SPI-2 (*S*.Tm^ΔSPI-2^)-encoded T3SS for 4 days. We compared *S.* Tm loads and microbiota composition to *S*. Tm^WT^ and S. Tm^avir^ data shown in Fig. 1. *S*. Tm^ΔSPI-1^ and *S*. Tm^ΔSPI-2^ colonized the intestine at largely similar levels (**Fig. S3A**). Interestingly, systemic *S*. Tm loads were similar between the mutants (**Fig. S4ABC**). *S*. Tm^ΔSPI-2^ (triggering gut inflammation via the SPI-1 T3SS) induced more cecal pathology and higher *S*. Tm abundance compared to the SPI-1 deficient mutant, although this difference was not statistically significant (**Fig. S3BC**). Microbiota composition in cecum content was significantly different between all groups, as determined by ADONIS Permanova analysis on the Bray–Curtis distance metric (**Fig. S3E, Table S3**). Blooms were only evident in *S*.Tm^ΔSPI-2^ and *S*. Tm^WT^ infected mice (**Fig. S3F**). Total bacterial load in *S*.T m^ΔSPI-1^ and *S*. Tm^ΔSPI-2^ infected mice was altered when compared to *S*. Tm^avir^ but not significantly changed when compared to the S. Tm^WT^ infected group (**Fig. S3G).** This applied also for most of the individual OMM^12^ strains (**Fig S4D-O**). Taken together, *S*. Tm^WT^ induced strong gut inflammation in OMM^12^ mice, beginning at day 3 p.i. and depending on the activity of both *Salmonella* type three secretion systems (T3SS). Severe inflammation at day 4 p.i. was paralleled by *S*. Tm^WT^ blooming, drastic reduction of total gut luminal bacterial loads and overall changes in microbiota composition.

### Anaerobic respiration and ethanolamine utilization fuels *S*. Tm blooms and dysbiosis

To analyze the relevance of anaerobic and aerobic respiration for *S*. Tm blooming, we generated *S*. Tm mutants deficient in nitrate respiration (*S*. Tm^Ni.;^ Δ*narZ*; *narG*::*cat*; *napA*::*aphT*) and in nitrate and tetrathionate respiration (*S*. Tm^Ni. + Te.^; Δ*narZ*; *narG*::*cat*; *napA*::*aphT*; *ttrS*::*tet*). Moreover, to probe the role of siderophore-mediated iron acquisition and ethanolamine utilization, we generated a *S.* Tm *entA*::cat (*S.* Tm*^EntA^*) and *eutC*::*aphT* mutant strain (*S*. Tm^EA^). In addition, we created a *S.* Tm *cyxA* mutant, attenuated in aerobic respiration by deletion of the cytochrome bd-II oxidase gene (*S.* Tm^cyx^; Δ*cyxA*).

We infected groups of OMM^12^ mice with 5 x 10^7^ CFUs of the mutant strains or *S*. Tm^WT^. Fecal samples were collected at different days after infection in order to monitor the course of intestinal *S.* Tm colonization. Mice were sacrificed at day 4 p.i. and cecal content was harvested for microbiota analysis, inflammation was quantified and systemic *Salmonella* loads were determined (**Fig. 3**). *S.* Tm strains colonized the gut of OMM^12^ mice equally well (**Fig. 3A**). Only *S*. Tm^EA^ showed significantly reduced loads at day 3 p.i. compared to *S*. Tm^WT^ (p<0.01, Kruskal-Wallis test with Dunn’s multiple comparison test; **Fig. 3A**). There was no difference in systemic *Salmonella* loads at day 4 p.i. between groups (**Fig. S5**). *S*. Tm^WT^, *S*. Tm^EA^ and S. Tm*^EntA^* induced profound colitis by day 4 p.i., while *S*. Tm^Ni. + Te^ induced significantly less colitis symptoms (p<0.001, **Fig. 3B**). For *S.* Tm^cyx^, no significant differences could be detected in pathogen loads in intestine or organs in comparison to *S*. Tm^WT^ (**Fig. S6**). Permanova analysis with ADONIS on the Bray–Curtis distance metric revealed significant differences between cecal community structure of *S*. Tm^Ni. + Te^ and *S*. Tm^WT^ or S. Tm*^EntA^* infected mice (**Fig. 3D**; **Table S4**) and abundance of most OMM^12^ bacteria was increased in *S*. Tm^Ni. + Te^ infected mice (**Fig. S7**). For *S.* Tm^cyx^, no significant difference was found with respect to microbiota composition in comparison to *S*. Tm^WT^ (**Fig. S6, 57**; **Table S5**). *S*. Tm^Ni. + Te^ mutant did not establish “blooms”, while relative abundances of the other *S*. Tm mutants were similar to *S.* Tm^WT^ (**Fig. 3E**) and total bacterial loads were increased in mice infected with *S*. Tm^Ni. + Te^ compared to *S.* Tm^WT^ infected mice (**Fig. 3F**).

**Fig. 3:**
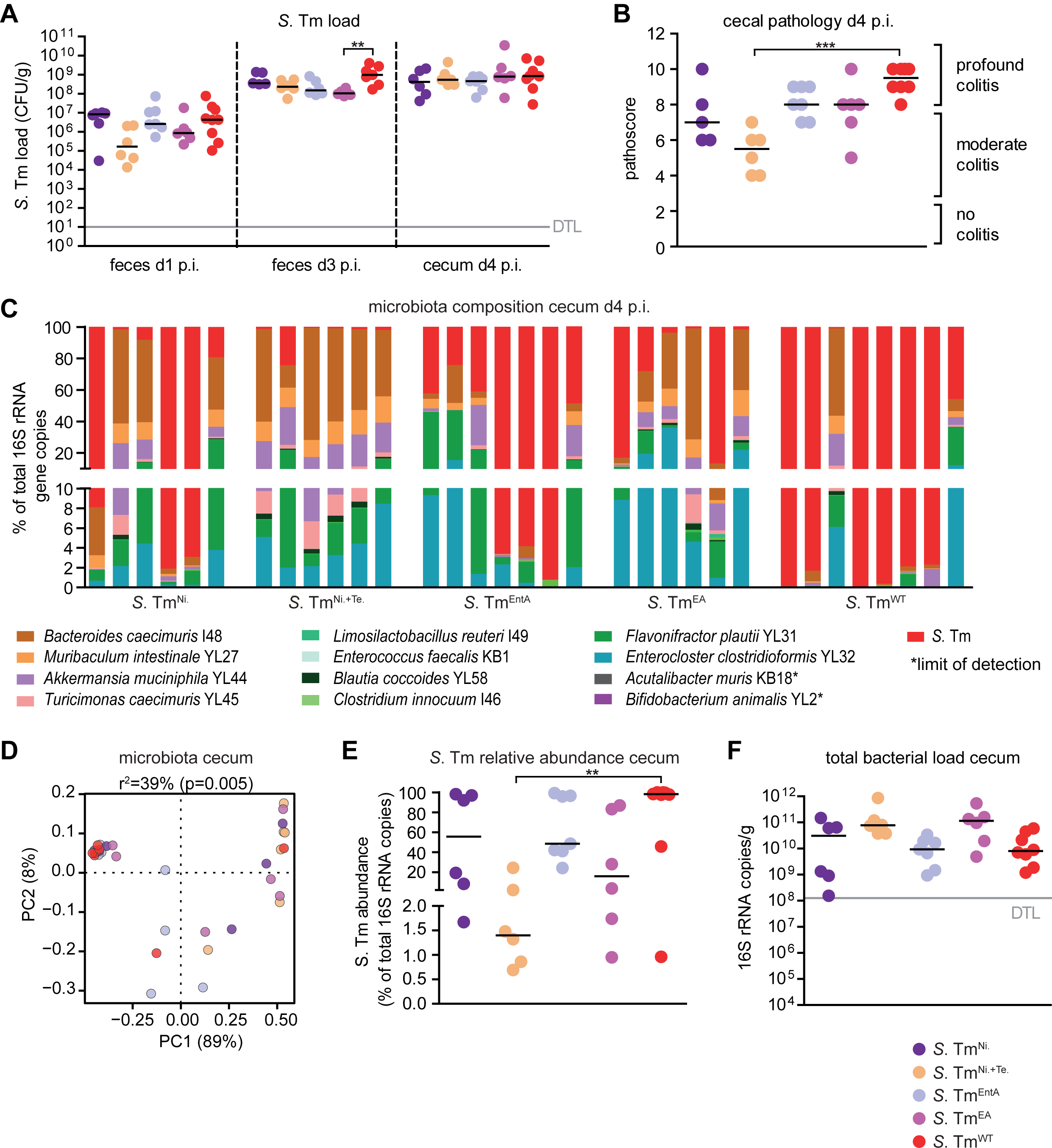
Course of infection of *S.* Tm mutants in anaerobic respiration, salmochelin production and ethanolamine utilization in OMM^12^ mice. OMM^12^ mice were orally infected with different *S.* Tm mutant strains: *S*. Tm^Ni.^ (n=6), *S*. Tm^Ni. + Te.^ (n=6), *S*. Tm^EntA^ (n=7), *S*. Tm^EA^ (n=6), or *S*. Tm^WT^ (n=8). Mice were sacrificed at day 4 post infection **(A)** *S.* Tm loads in feces and cecal content at days 1, 3 and 4 post infection were determined by plating. **(B)** Histopathological analysis of cecal tissue at day 4 p.i.. Cecal tissue sections of the mice were stained with hematoxylin/eosin to determine the degree of submucosal edema, neutrophil infiltration and epithelial damage (1–3: no pathological changes; 4–6: moderate inflammation; above 7: severe inflammation). **(C)** Analysis of microbiota composition in cecal content. Microbiota composition was determined by strain-specific qPCR assay and is shown as relative abundances of the individual strains (% of cumulated 16S rRNA gene copy numbers). **(D)** PCoA based on the distance matrix of Bray-Curtis dissimilarity of relative OMM^12^ abundance profiles shows the effect of the different *S*. Tm mutant strains. Points are colored by time (days) after infection. **(E)** Relative abundance of *S*. Tm and **(F)** absolute amount of 16S rRNA gene copies (determined by an universal primer / probe combination) in cecal contents 4 days p.i.. Statistical analysis was performed using Kruskal-Wallis test with Dunn’s multiple comparison test (* p<0.05, ** p<0.01, *** p<0.001). Each dot represents one mouse, black lines indicate median, grey lines indicate detection limit (DTL).

### Neutrophils reduce overall luminal bacterial loads but do not promote *Salmonella* ‘blooms’

Next, we set out to test the hypothesis that neutrophils are the cause of decreased luminal bacterial loads in S. Tm^WT^ infected OMM^12^ mice at day 4 p.i.. Therefore, we employed a neutrophil depletion protocol using α-Ly6G and α-G-CSF antibodies or isotype control antibodies in OMM^12^ mice that were additionally infected with *S*. Tm^WT^ (**Fig. 4A**). Antibody mediated depletion was confirmed by FACS analysis from blood samples gating on CD45^+^, SYTOX^-^, CD3^-^, CD11b^+^, Ly-6G^+^, Ly-6C^intermediate^ cells and by quantifying neutrophil-elastase in cecal content (**Fig. 4B**). Only mice with confirmed neutrophil depletion (< 4% Ly6G^+^/CD45^+^ cells) were included in the analysis.

**Fig. 4:**
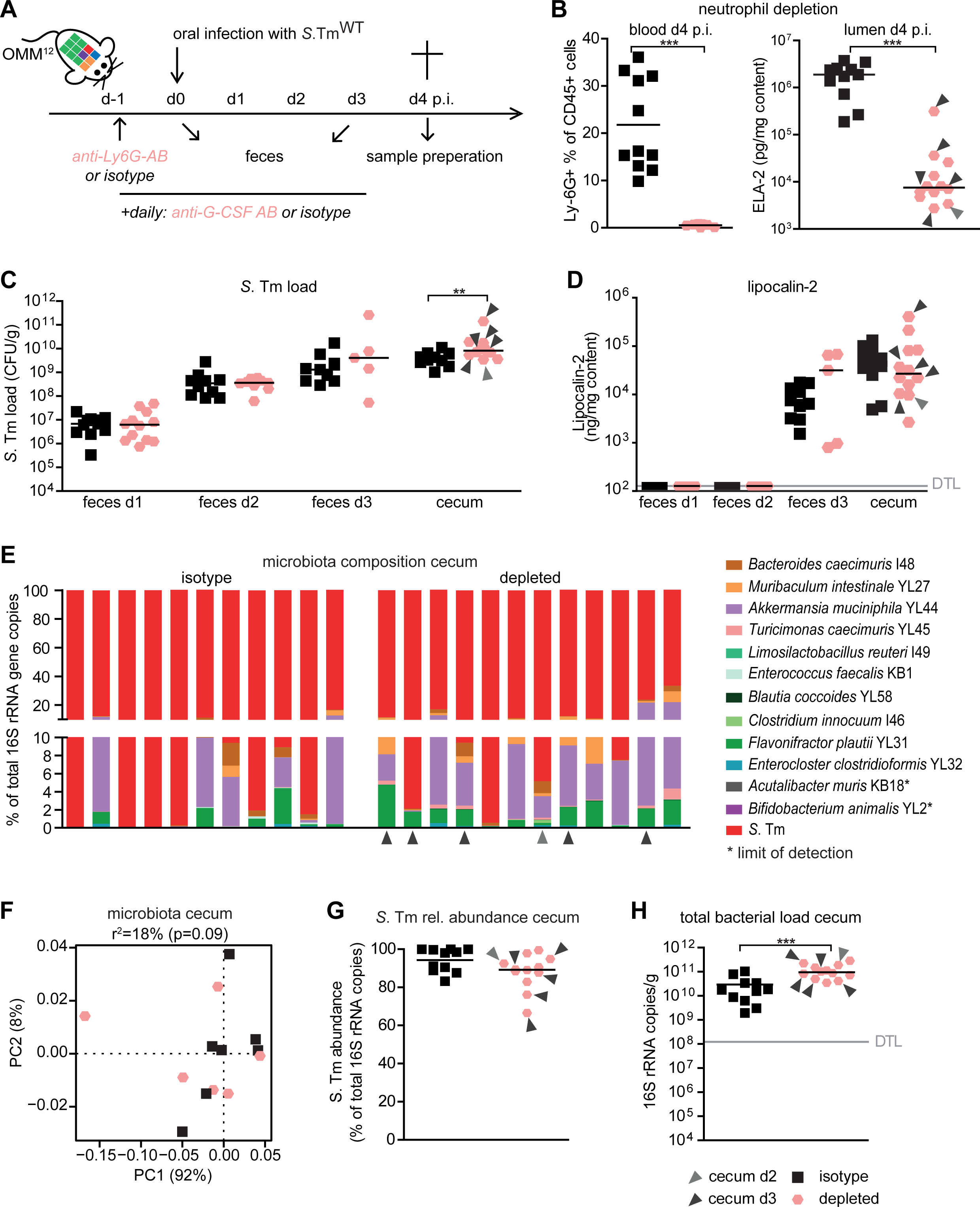
Neutrophils reduce total gut luminal bacterial loads but are dispensable for the induction of *Salmonella* ‘blooms’ and dysbiosis. **(A)** Experimental scheme. OMM^12^ were treated with one dose of α-Ly6G antibody or isotype control one day before infection with *S*. Tm^WT^. In addition, α-G-CSF antibody or isotype control were daily administered starting from day −1 until day 3 p.i.. All antibodies were given via the intraperitoneal route. **(B)** Monitoring of neutrophil depletion at day 4 p.i. in blood by FACS gating on CD45^+^, SYTOX^-^, CD3^-^, CD11b^+^, Ly-6G^+^, Ly-6C^- intermediate^ cells or in cecal content by neutrophil elastase specific ELISA. FACS data are indicated as the mean. Statistical analysis was performed using Mann Whitney test. **(C)** *S.* Tm loads in feces at days 1, 2 and 3 and cecal content at day 4 p.i. were determined by plating. **(D)** Lipocalin-2 levels in feces and cecal content (ng / mg content). Lipocalin-2, elastase levels and CFUs are shown as median and data were analyzed using Mann Whitney test. **(E)** Analysis of cecal microbiota composition at day 4 p.i. with different *S.* Tm^WT^ and neutrophil depletion or isotype control treatment. Microbiota composition is shown as relative abundance and expressed as % of cumulated 16S rRNA gene copy numbers. **(F)** PCoA based on the distance matrix of Bray-Curtis dissimilarity of relative OMM^12^ abundance profiles shows the effect of the different treatments. **(G)** Relative abundance of *S*. Tm and **(H)** absolute amount of 16S rRNA gene copies (determined by an universal primer / probe combination) in cecal contents 4 days p.i.. Light and dark gray arrows indicate samples analyzed at day 2 or 3 p.i., respectively. Statistical analysis was performed using Mann Whitney test (* p<0.05, ** p<0.01, *** p<0.001). Each dot represents one mouse, black lines indicate median, grey lines indicate detection limit (DTL).

Some neutrophil-depleted mice developed severe infection and had to be euthanized at day 2 or 3 p.i. (indicated with gray and black arrows). Cecal *S*. Tm^WT^ loads were similar in both groups of mice at day 1-3 p.i. but significantly higher in neutrophil-depleted mice at day 4 p.i. (**Fig. 4C**). At the point of sacrifice, all mice developed pronounced gut inflammation as determined by LCN-2 levels (**Fig. 4D**). Systemic loads of *S*. Tm^WT^ were higher in the mLN, spleen and liver of neutrophil-depleted mice (**Fig. S8A-C**).

There was no apparent difference in the overall cecal microbiota composition between the groups at day 4 p.i. (**Fig. 4EF**; **Table S6**). Moreover, no difference in rel. *S.* Tm^WT^ abundance was observed between the groups (**Fig. 4G**). However, a lower overall amount of total gut luminal bacteria was observed in the isotype group (**Fig. 4H** copies/mg content isotype vs. depleted p<0.001, Mann Whitney test), indicating that neutrophils are causal for inflammation-associated reduction of gut luminal bacterial loads in *S.* Tm infected OMM^12^ mice. Reduction was most pronounced for *Bacteroides caecimuris* I48, *Muribaculum intestinale* YL27, *Akkermansia muciniphila* YL44, *Turicimonas muris* YL45, *Blautia coccoides* YL58, *Flavonifractor plautii* YL31 and *Enterocloster clostridioformis* YL32 – other bacteria were less affected (**Fig. S8D-O**).

## Discussion

Together, the host and its intestinal symbionts establish microbiota-nourishing immunity that forms an efficient barrier against *S.* Tm infection (42). Epithelial hypoxia, short-chain fatty acid production and depletion of growth substrates are cornerstones of this barrier. Using its virulence factors, *S*. Tm can overcome colonization resistance, trigger inflammation and alter the gut luminal environment to benefit from it – at the expense of intestinal commensal bacteria. Inflammation-driven pathogen “blooming” has a major impact on *S.* Tm dissemination and evolution (43). Overgrowth of related Enterobacteriaceae promotes exchange of conjugative plasmids (44) and phages (45). This increases the chance for spreading antibiotic resistances and fosters evolution of highly virulent strains (46), emphasizing the broad implications of S. Tm adaptation of the inflammatory milieu in the mammalian gut. Here we analyze the extent to which *S.* Tm-induced intestinal inflammation, virulence and gut luminal neutrophils contribute to alterations in the gut microbiota loads and community profiles in OMM^12^ mice, a widely used infection model (4, 9, 47) based on a well-characterized minimal bacterial community (48).

Wild type *S*. Tm induced severe gut inflammation within 4 days after oral infection of OMM^12^ mice, which was characterized by an overall reduction of luminal bacterial loads and *S*. Tm blooming (i.e. *S.* Tm being the dominant species). This overgrowth was not observed with an avirulent mutant (Δ*invG* Δ*ssaV*; *S.* Tm^avir^), which can colonize the gut but is defective in inducing inflammation. This confirms results from previous studies in the streptomycin-treated mouse colitis model and highlights the pronounced negative impact of intestinal inflammation on the majority of intestinal commensals while fostering pathogen overgrowth (15).

Comparing the course of infection in *S*. Tm TTSS mutants, we found that *S*. Tm^ΔSPI-1^ was more strongly attenuated in inducing gut inflammation than *S*. Tm^ΔSPI-2^ at day 4 p.i.. This suggests that in OMM^12^ mice, the SPI-1 TTSS contributes more to overall inflammation than the SPI-1 TTSS. Previous work in the streptomycin model showed that SPI-1 triggers inflammation within hours after infection. This response is due to effector proteins that initiate a complex immunological signaling cascade, culminating in the production of IFNγ (49–53). In contrast, in OMM^12^ mice, overt inflammation starts only by day 3 p.i. (**Fig. 1FG**), which is mainly due to initially lower *S.* Tm loads (<10^7^ CFU/g) compared to streptomycin-treated mice (>10^8^ CFU/g). Mutants lacking a functional SPI-1 or a SPI-2 TTSS caused equally attenuated colitis at day 4 p.i.. However, at early time points post infection (day 1, 2 p.i.) a *S*. Tm^ΔSPI-1^ mutant is strongly attenuated and inflammation triggered via the SPI-2 TTSS only developed at day 3 p.i. (54).

*S.* Tm mutants in anaerobic respiration as well as iron uptake via salmochelin and ethanolamine utilization were previously shown to be attenuated in growth in the inflamed gut in direct competition with *S.* Tm^WT^ (11, 20, 27, 55). In this study, we aimed to identify mechanisms that promote *S.* Tm blooming in competition against the OMM^12^ microbiota. We show that a double mutant in anaerobic nitrate and tetrathionate respiration (*S*. Tm^Ni+Te^) has a reduced capacity to bloom and induce microbial community shifts. As gut inflammation in *S*. Tm^Ni+Te^ infected mice was also significantly reduced at day 4 p.i., we cannot conclude that anaerobic respiration creates an advantage in competition against OMM^12^ in the inflamed gut. In contrast, mutants in O_2_ and nitrate respiration, ethanolamine utilization and salmochelin production (*S.* Tm^cyx^*, S*. Tm^Ni^, *S*. Tm^EA^, *S.* Tm*^EntA^*) induced strong inflammation, bloomed and altered microbiota composition to a similar extend as *S.* Tm^WT^. Therefore, these mechanisms are, at least individually, not required for competition against OMM^12^ bacteria in the inflamed gut. In previous work using the streptomycin-treated colitis model, *S.* Tm^cyx^ was strongly attenuated in intestinal colonization (10). This difference might be due to the presence of a complex microbiota including other facultative anaerobic bacteria that compete with *S.* Tm for nutrients and O_2_ in the inflamed gut.

These data indicate that neutrophils protect the host from systemic Salmonella and reduce total gut luminal bacterial loads but do not contribute to *Salmonella* ‘blooms’ and concomitant dysbiosis. Notably, antibody-mediated depletion of neutrophils resulted in reduction of lipocalin-2 and elastase-2 levels and concomitantly resulted in an overall increase in total gut luminal bacterial loads. This confirms the pronounced antibacterial function of transmigrated neutrophils in the gut lumen reported previously (34, 35, 37). In contrast, neutrophil depletion did not change the relative abundance of gut luminal *S.* Tm. This suggests that neutrophil-derived antimicrobial effectors equally reduce numbers of *S.* Tm and members of the OMM^12^ and are not causally involved in *S.* Tm blooming in this model. Rather, “blooming” is mainly mediated by the pathogens’ diverse capabilities of taking advantage of the altered substrate range in the inflamed intestine.

Gnotobiotic OMM^12^ mice are increasingly used as standardized infection model for *S.* Tm and other pathogenic Enterobacteriaceae (4, 9, 41). Here, we present detailed characterization of the course of *S.* Tm^WT^ infection and mutants in major virulence factors. Our study reveals that OMM^12^ mice infected with *S.* Tm^WT^ develop fulminant colitis by day 4 p.i. characterized by pathogen blooming and shifts in microbiota composition. We conclude that OMM^12^ mice are a valuable model to dissect the impact of *S.* Tm^WT^ induced inflammation on the gut ecosystem including all its members.

## Materials and Methods

### Bacterial strains

For detailed information about individual Oligo-MM^12^ strains refer to (40). All Oligo-MM^12^ strains are part of the mouse intestinal bacteria collection (miBC (56)) and are also available via the German Collection of Microorganisms and Cell Cultures (www.dsmz.de/miBC). *Salmonella* strains used and created in this study are based on *Salmonella enterica* serovar Typhimurium strain SL1344 (Table 1).

**Table 1:**
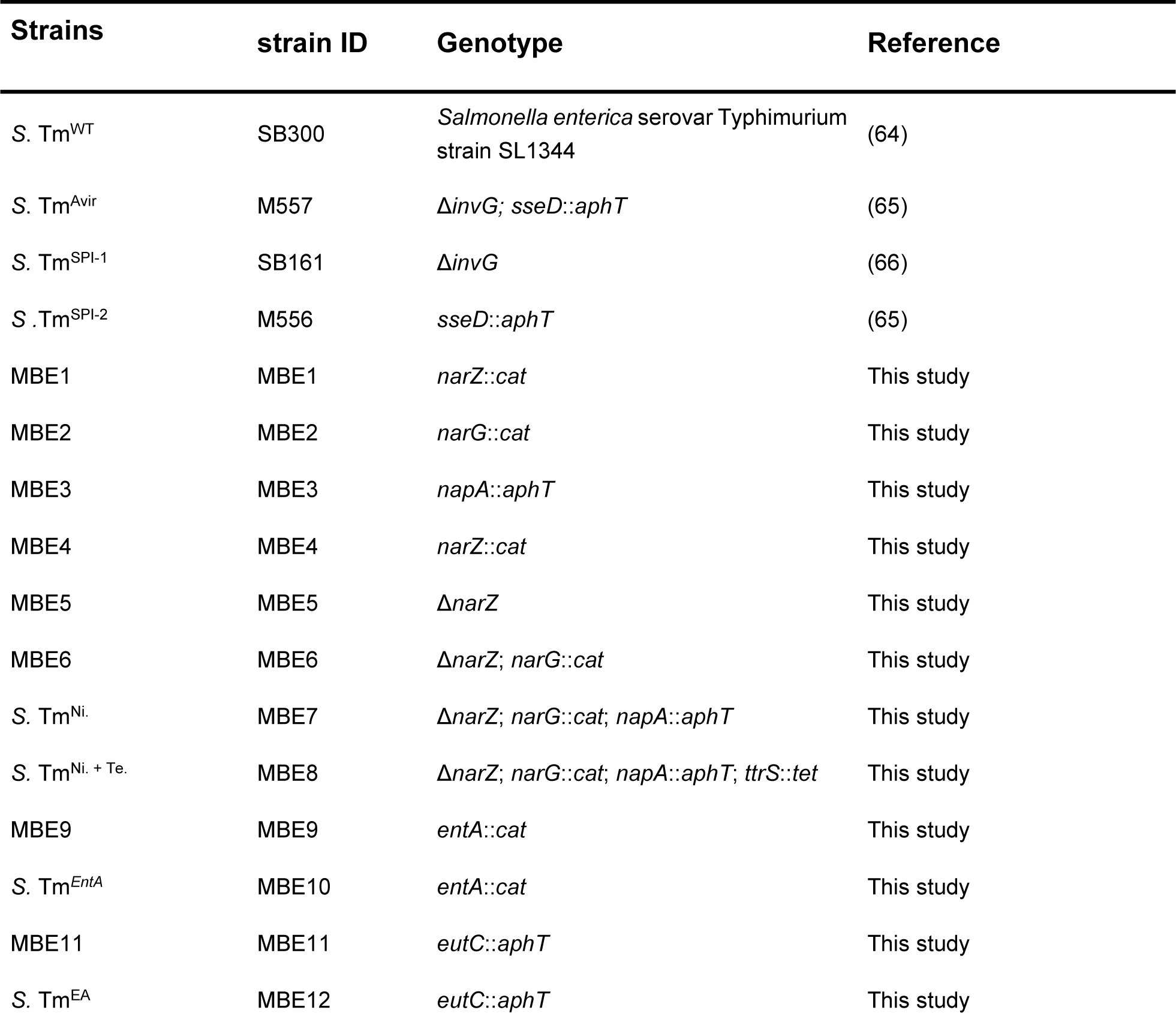
Bacterial strains used in this study.

| Strains | strain ID | Genotype | Reference |
| --- | --- | --- | --- |
| *S. Tm*<sup>WT</sup> | SB300 | *Salmonella enterica* serovar Typhimurium strain SL1344 | (64) |
| *S. Tm*<sup>Avir</sup> | M557 | $\Delta invG$; *sseD::aphT* | (65) |
| *S. Tm*<sup>SPI-1</sup> | SB161 | $\Delta invG$ | (66) |
| *S. Tm*<sup>SPI-2</sup> | M556 | *sseD::aphT* | (65) |
| MBE1 | MBE1 | *narZ::cat* | This study |
| MBE2 | MBE2 | *narG::cat* | This study |
| MBE3 | MBE3 | *napA::aphT* | This study |
| MBE4 | MBE4 | *narZ::cat* | This study |
| MBE5 | MBE5 | $\Delta narZ$ | This study |
| MBE6 | MBE6 | $\Delta narZ$; *narG::cat* | This study |
| *S. Tm*<sup>Ni.</sup> | MBE7 | $\Delta narZ$; *narG::cat*; *napA::aphT* | This study |
| *S. Tm*<sup>Ni. + Te.</sup> | MBE8 | $\Delta narZ$; *narG::cat*; *napA::aphT*; *ttrS::tet* | This study |
| MBE9 | MBE9 | *entA::cat* | This study |
| *S. Tm*<sup>EntA</sup> | MBE10 | *entA::cat* | This study |
| MBE11 | MBE11 | *eutC::aphT* | This study |
| *S. Tm*<sup>EA</sup> | MBE12 | *eutC::aphT* | This study |

### Construction of *S*. Tm strains and plasmids

All bacterial strains used in this study are listed in Table 1. ***S*. Tm^Ni^**. (MBE7: Δ*narZ*, *narG*::*cat*, *napA*::*aphT*) was generated in a step wise manner. First, single *S*. Tm mutant strains were generated (MBE1: *narZ*::*cat*, MBE2: *narG*::*cat* and MBE3: *napA*::*aphT*) using λ red recombination as described previously (57). Briefly, antibiotic resistance genes as well as FRT-sites were PCR amplified using specific knock out primers (ko-primers, Table 2) and the plasmids pKD3 (*cat*) or pKD4 (*aphT*) listed in Table 3. PCR products were subsequently electroporated into *S*. Tm^WT^ harboring pKD46 (Table 3). The *narZ*::*cat* allele from MBE1 was transduced to *S*. Tm^WT^ by P22-transduction (58) in order to create MBE4. The *narZ*::*cat* was then deleted using FLP-recombinase encoded on pCP20 (Table 3). The resulting strain MBE5 (Δ*narZ*) was transduced with P22-phage lysate of MBE2 in order to create MBE6 (Δ*narZ*, *narG*::*cat*). MBE6 was finally transduced with P22-phage lysate of MBE3 to generate MBE7 (Δ*narZ*, *narG*::*cat*, *napA*::*aphT*). In order to construct ***S*. Tm^Ni. + Te.^** (MBE8: Δ*narZ*, *narG*::*cat*, *napA*::*aphT*, *ttrS*::*tet*), the *ttrS*::*tet* allele from *S*. Tm stain M961 (Δ*sodCI*, Δ*sodCII*, *BCB4*::*tet*, (59)) was P22-transduced into MBE7. ***S*. Tm^EntA^** (MBE10: *entA*::*cat*) was created by P22-transduction of the *entA*::*cat* allele from MBE9 to *S*. Tm^WT^. MBE9 was previously generated by λ red recombination. ***S*. Tm^EA.^** (MBE12: *eutC*::*aphT*) was constructed by P22-transduction of the *eutC*::*aphT* allele from MBE11 to *S*. Tm^WT^. MBE11 was generated before using λ red recombination. In all cases, genetic modifications were verified by PCR using gene specific check-up primers (Table 2). The construction and sequences of plasmids harboring full length 16S rRNA sequences used as standard for qPCR reactions are detailed in (40).

**Table 2:**
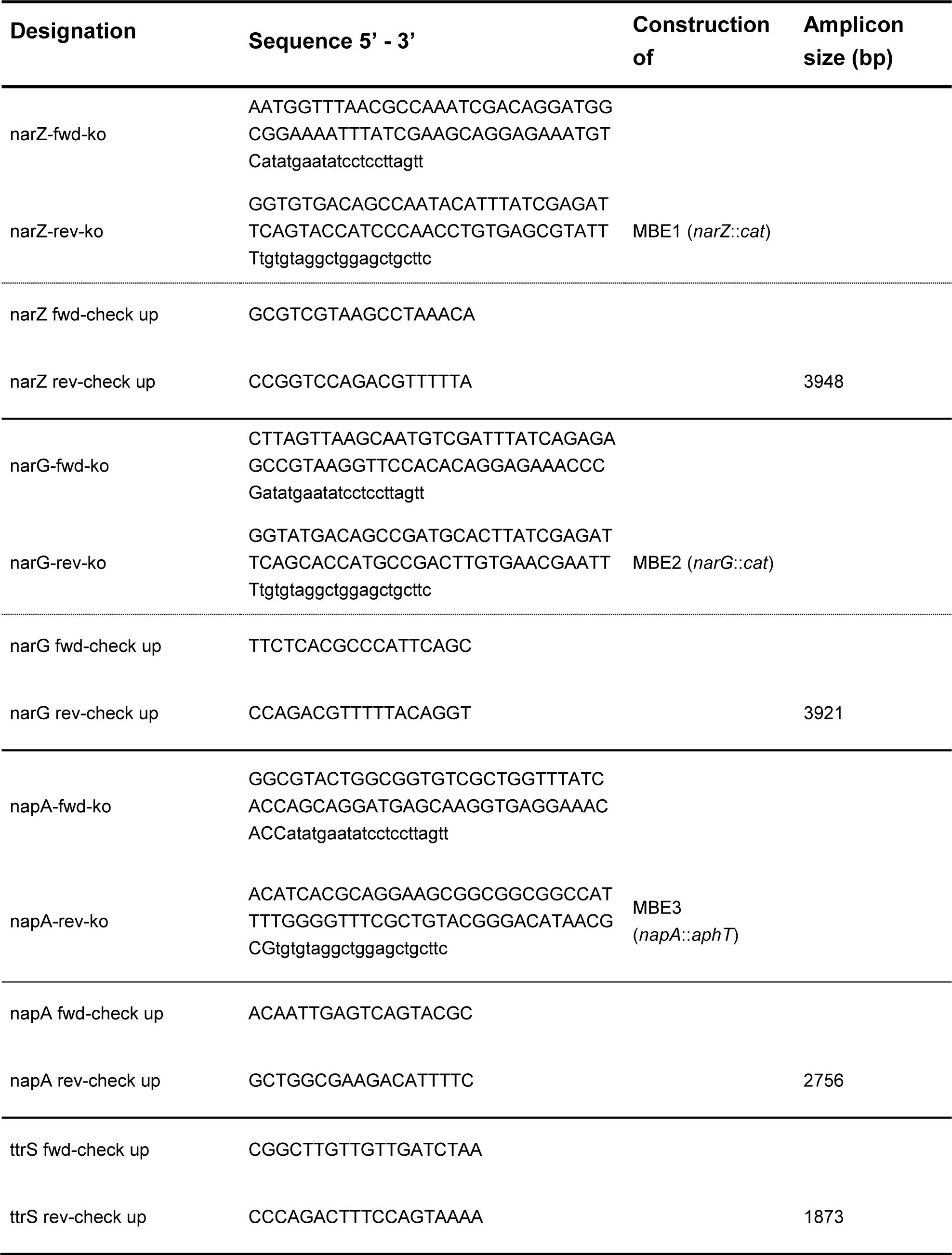
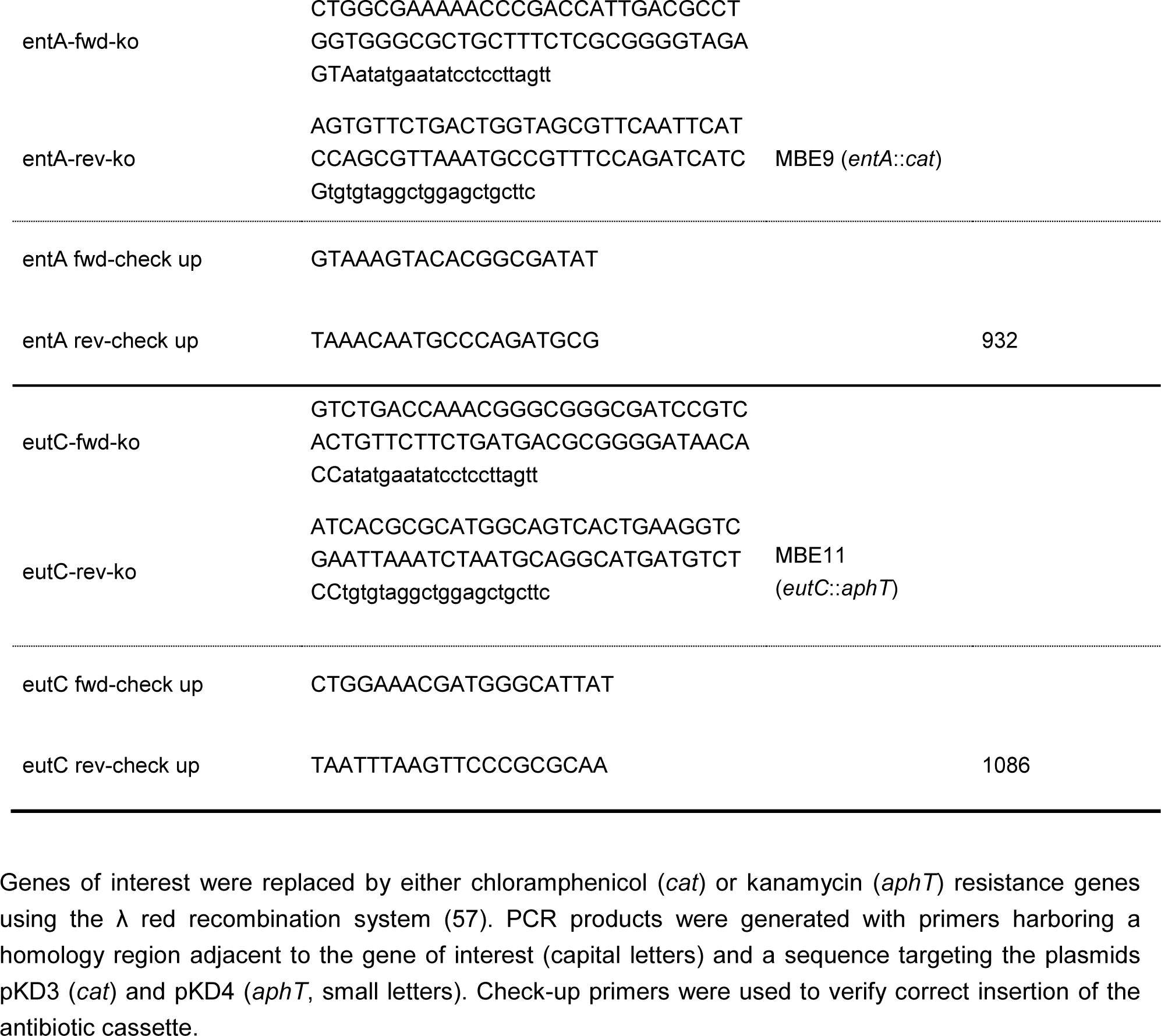
Primers used for construction of *Salmonella* mutants.

**Table 3:** Plasmids used for λ red and Flp recombination.

| Plasmid | Function | Genotype | Reference |
| --- | --- | --- | --- |
| pKD3 | Harbors a chloramphenicol resistance gene (*cat*) | *cat* with FRT sites, Amp<sup>R</sup> | (57) |
| pKD4 | Harbors a kanamycin resistance gene (*aphT*) | *aphT* with FRT sites, Amp<sup>R</sup> | (57) |
| pKD46 | Harbors 2,154 nt (31088–33241) of phage $\lambda$, enzymes for recombination | *araC*, ParaB $\gamma$, $\beta$ and *exo*, *repA101*<sup>ts</sup>, *Amp*<sup>R</sup> | (57) |
| pCP20 | Harbors FLP-recombinase | *FLP*<sup>+</sup>, $\lambda$ *cl857*<sup>+</sup>, $\lambda$ *p<sub>R</sub>* *Rep*<sup>ts</sup>, *Amp*<sup>R</sup>, *Cm*<sup>R</sup> | (67) |

### Culture conditions and bacterial *in vitro* assays

Culture conditions as well as cryopreservation protocols of individual Oligo-MM^12^ strains are described in (40). For mouse infection, a single colony of a *S*. Tm strain was inoculated in 3 ml LB medium containing 0.3 M NaCl (LB^0.3^) and grown on a wheel rotor for 12 h at 37°C. The starter culture was diluted 1:20 in fresh LB^0.3^ and incubated for another 4 h at 37°C under constant rotation. The subculture was washed in ice-cold PBS and the pellet was subsequently re-suspended in PBS. Mice were finally gavaged with 5 x 10^7^ CFUs of *S*. Tm strains. For investigating phenotypes conferred by gene knock outs, individual colonies of *S*. Tm strains were aerobically inoculated with 5 ml M9 medium (Na_2_HPO_4_ · 2H_2_O 40 mM, KH_2_PO_4_ 20 mM, NaCl 9 mM, NH_4_Cl 37.39 mM, D-glucose 1.1 mM, MgSO_4_ 1 mM, CaCl_2_ 100 μM, Thiamine 10 mg/ml, Histidine 500 mg/l) without antibiotics on a wheel rotor for 12 h at 37°. Aerobic as well as anaerobic (7% H_2_, 10% CO_2_, rest N_2_) subcultures with a starting OD_600_ of 0.02 were subsequently carried out in 10 ml M9 medium with and without nitrate (20 mM). For testing *S*. Tm strains deficient in ethanolamine utilization, M9 medium was additionally supplemented with ethanolamine (5 mM) and tetrathionate (40 mM). Aerobic subcultures were performed in glass Erlenmeyer flaks sealed with aluminum foil, whereas anaerobic subcultures were conducted in 100 ml Wheaton glass serum bottles with pre-reduced medium (pre-reduction for at least 2 days under anoxic conditions: 3% H_2_, rest N_2_). Subcultures were incubated at 37°C, shaken at 180 rpm and samples were taken for OD_600_ measurement and determination of nitrate concentrations (60). *In vitro* competition assays were performed as described in (61). Briefly, starter cultures of the competing strains in 5 ml LB or M9 medium without antibiotics incubated on a wheel rotor for 12 h at 37°C were mixed in a 1:1 ratio with a starting OD_600_ of 0.05 of each strain and used to inoculate 14 ml cultures with either plain mucin broth (Type II porcine mucin 2.5 g/l, Morpholino propanesulfonic acid (MOPS) sodium 9.25 g/l, MgSO_4_ 2.2 mM, trace elements 1000x: CaCl_2_ · 2H_2_O 0.3 mM, ZnSO_4_ · 7H_2_O 0.1 mM, FeSO_4_ · 7H_2_O 0.045 mM, Na_2_SeO_3_ 0.2 mM, Na_2_MoO_4_ · 2H_2_O 0.2 mM, MnSO_4_ · H_2_O 2 mM, CuSO_4_ · 5H_2_O 0.1 mM, CoCl_2_ · 6H_2_O 3 mM, NiSO_4_ · 6H_2_O 0.1 mM) or mucin broth containing nitrate (40 mM). For competition assays underlining the importance of tetrathionate respiration, the mucin broth was supplemented with tetrathionate (40 mM) and histidine (500 mg/l) to enhance growth. Cultures were statically incubated in 15 ml plastic tubes for 24 h at 37°C. CFUs of *S*. Tm strains were determined by plating on LB agar plates containing selective antibiotics. The deficiency of acquiring iron via siderophores was investigated using Chromazurol S (CAS) agar. 1 ml of starter cultures in 5 ml LB medium without antibiotics incubated on a wheel rotor for 12 h at 37°C was centrifuged and the pellet was resuspended in sterile ice-cold PBS. 5 µl were adjusted to an OD_600_ of 0.1, spotted on CAS agar plates and incubated at 37°C for 48 h. The diameters of the colony as well as the diameter of the colony + the orange CAS halo were measured and the CAS-reactive ring was calculated in mm according to the formula: 0.5 x [(diameter of colony + CAS halo) – diameter of colony].

### DNA extraction from intestinal contents

Fecal and cecal DNA were extracted using the QIAamp DNA Stool Mini Kit (Qiagen) according to the manufacturer’s protocol with modifications. The protocol was extended with an initial bead-beating step using glass beads (0.5-0.75 mm) and zrikonia beads (< 100 µm). Lysozyme (20 mg/ml) was additionally added to the lysis buffer.

### Quantitative PCR (qPCR) of 165 rRNA genes

16S rRNA specific primers and hydrolysis probes were designed and validated according to the MIQE guidelines as detailed previously (40, 62). qPCR reactions were performed in a LightCycler96 thermo cycler (Roche) using white LightCycler480 Multiwell Plate 96 plates (Roche). One 20 µl reaction contained 250 nM of hydrolysis probe, 300 nM of each corresponding primer, 5 ng of template gDNA, water and FastStart Essential DNA Probes Master (Roche). The sequences of primers and hydrolysis probes, the qPCR efficiencies as well as the specificity/detection limits of qPCR assays are depicted in (40). All qPCR reactions were performed in duplicates applying the following conditions: 95°C for 10min, followed by 45 cycles of 95°C for 15s and 60°C for 1min. Fluorescence for each cycle was recorded after the step at 60°C. Quantification cycle (Cq) as well as the baseline were automatically determined by the software LightCycler96 version 1.1 (Roche).

### Animal experiments

Oligo-MM^12^ mice were generated as previously described (40) and sterilely bred in the animal housing facility of the Max von Pettenkofer-Institute. Sex- and age-matched mice were orally infected with 5 x 10^7^ CFUs of *S.* Tm strains under germ-free conditions and maintained in gnotocages (Han, Bioscape). The protocol for neutrophil depletion was adapted from (36). Briefly, neutrophils were depleted by one i.p. injection with 150 µg of α-m Ly-6G (clone: 1A8, BioXCell) prior to infection with *S*. Tm^WT^ followed by daily i.p. injections with 10 µg of α-mouse G-CSF (clone: 67604, R&D Systems). Control mice received 150 µg of Rat IgG2a (clone: 2A3, BioXCell) once and daily injections with 10 µg of Rat IgG1 (clone: 43414, R&D Systems). Antibodies were dissolved in sterile PBS. Mice were euthanized by cervical dislocation and organs were aseptically removed. *Salmonella* loads in mesenteric lymphnodes, liver, spleen as well as in cecal content and feces were determined by plating on MacConkey agar plates (Roth). The mouse lipocalin-2/NGAL detection kit (R&D Systems) ELISA kit and HRP-Streptavidin (Biolegend) were used for measuring lipocalin-2 levels. Neutrophil specific elastase was quantified using the Mouse Neutrophil Elastase/ELA2 DuoSet ELISA kit and the DuoSet Ancillary Reagent Kit2 (both from R&D Systems). Cecal pathology was scored at necropsy. Optimal cutting temperature (OCT) compound (Sakura, Torrance) was used to embed cecal tissue. Tissues were flash frozen and cut in 5 µm cryosections. Cryosections were H&E-stained and scored as detailed in (63). Scoring parameters include: submucosal oedema, infiltration, loss of goblet cells as well as epithelial damage. Scoring was performed in a blinded manner and yielded in a total score of 0-13 points according to the severity of inflammatory symptoms (scores: 0 – 3: no pathological change, 4 – 8: mild inflammation, 9 – 13: severe inflammation). All animal experiments were approved by the local authorities (Regierung von Oberbayern) and were performed according to the legal requirements.

### FAC5

Efficient depletion of neutrophil in the blood was monitored by FACS. 3 - 4 drops of blood taken from the tail vein were collected in 1 ml pre-cooled FACS buffer (PBS and 1% heat inactivated FCS). For subsequent erythrolysis, samples were centrifuged and FACS buffer was exchanged by 1 ml of BD FACS^TM^ Lysing Solution (BD). After 10 min incubation in the dark at RT, samples were centrifuged again, Lysing Solution was discarded and cells were resuspended in 100 µl of FACS buffer. Cells were subsequently stained with α-CD45-PerCP (clone: 30-F11, Biolegend, 1:100), α-CD11b-APC-Cy7 (clone: M1/70, Biolegend, 1:200), α-Ly-6G-Pacific Blue (clone: 1A8, Biolegend, 1:200), α-Ly-6C-FITC (clone: AL-21, BD, 1:400), α-CD3-PE (clone: 17A2, Biolegend, 1:200) and α-CD16/CD32 (93, FC-Block, eBioscience, 1:100). Following 30 min of incubation at 4°C, cells were washed and analyzed with the FACSCANTO II (BD). SYTOX red (5 nM) was used to discriminate dead and living cells. Data were recorded using the BD FACSDivaTM software (BD) and analyzed using the FlowJo software. Neutrophils were identified as CD45^+^, SYTOX^-^, CD3^-^, CD11b^+^, Ly-6G^+^, Ly-6C^-intermediate^ cells.

### Statistical analysis

CFU data, LCN-2 levels and pathoscores were expressed as median. More than two different groups were compared to each other using Kruskal-Wallis test, with Dunn’s multiple comparison test. Two groups were compared using the Mann Whitney test (Prism 5; GraphPad Software, San Diego, CA, USA). The percentage of individual bacteria was expressed as mean +/-standard deviation (SD). Differences between individual bacteria were compared using a two-way ANOVA, with Bonferroni posttest (Prism 5; GraphPad Software, San Diego, CA, USA). FACS data were expressed as the mean and analyzed using unpaired t test (Prism 5; GraphPad Software, San Diego, CA, USA).

To analyze clustering of qPCR data, Pearson distance matrix was used containing dissimilarity values for each pairwise comparison. Strength and statistical significance of sample grouping were determined applying the nonparametric Adonis method based on the permutational multivariate ANOVA (PERMANOVA), together with the parametric significance test PERMDISP, which analyzes multivariate homogeneity of group dispersions. The used scripts are available in QIIME (Caporaso et al. 2010).

In all cases, p values < 0.05 were considered as statistically significant.

Fold changes in absolute abundance were calculated with absolute values that were normalized to a million gene copies determined by universal probe.

## Acknowledgments.

The work was supported by grants from the BMBF (Medizinische Infektionsgenomik), DFG Priority program SPP1656, research grants DFG STE 1971/2-1, the German Center for Infection Research (DZIF). All authors declare they have no competing financial or intellectual interests. M.B. and B.S. conceived and designed the study. M.B., C.E. and B.S. and wrote the paper. M.B., C.E., S.H., D.R., T.D., S.B., P.S., S.H., M.D. performed the experiments. M.B., C.E., D.G., P.M., T.D. analyzed the data. M.B. and A.B. contributed important material. All authors approved the definitive version of the manuscript.

